# The impact of distillery effluent irrigation on plant-growth-promoting traits and taxonomic composition of bacterial communities in agricultural soil

**DOI:** 10.1101/554709

**Authors:** Priyanka Kumari, Binu M. Tripathi, Ram N. Singh, Anil K. Saxena, Rajeev Kaushik

## Abstract

Long-term irrigation of agricultural fields with distillery effluent (DE) may alter the physical, chemical and biological properties of the soil. Microorganisms are critical to the maintenance of soil health and productivity. However, the impact of DE irrigation on activity and taxonomy of soil microorganisms is poorly understood. Here we studied plant-growth-promoting (PGP) traits and taxonomic composition of bacterial communities in agricultural soil irrigated with DE in conjugation with irrigation water, using cultivation-dependent and - independent methods. Most of the bacterial isolates obtained from DE irrigated soil were found to display PGP traits (phosphate solubilization, siderophore, indolic compounds and ammonia production). Diverse bacterial taxa were found in both culturable bacterial community and 16S rRNA gene clone library, which belonged to bacterial phyla *Proteobacteria* (Alpha-, Beta- and Gamma- subdivisions), *Firmicutes, Actinobacteria, Acidobacteria, Bacteroidetes* and *Gemmatimonadates*. Overall, these results indicate that PGP traits and taxonomic diversity of soil bacterial communities were not severely impacted by DE irrigation.

## Introduction

Partially treated or untreated anaerobically digested distillery effluent (DE) is mainly discharged in non-judicious manner on to the agricultural lands and in local waterways in India. Distillery effluent is of brown color with high salt levels and has high biological and chemical oxygen demands (BOD = 40,000-50,000 mg L^−1^ and COD = 80,000-100,000 mg L^−1^) [1]. In India, controlled and judicious application of DE in agriculture is considered as one of the viable option to transform industrial waste to value added resource as it contains considerable amounts of both macro- and micronutrients [2,3,4,5]. However, non-judicious application of DE has also been shown to adversely impact the growth and yield of crop plants [6,7].

In an agroecosystem, the nutrient accessibility and productivity is largely depends on soil microorganisms, as they play some vital functions such as nutrient cycling [8], soil development [9] and organic matter decomposition [10]. Several studies have shown that DE irrigation alters the soil physico-chemical properties [3,11,12], which, in turn, influence the activity and biomass of soil microorganisms [13,14]. However, the impact of DE irrigation on plant-growth-promoting (PGP) traits and taxonomic composition of soil microbial communities is still poorly understood. Two previous studies have shown that application of industrial effluent to agricultural soils could increase the diversity and catabolic profile of microbial communities [15,16]. However, these studies only considered pulp and paper mill effluent irrigation. Therefore, it is important to study the impact of DE irrigation on PGP traits and taxonomic composition of soil microbial communities.

It is now well documented that only a small proportion (1–5%) of the total soil microorganisms can be cultured on currently known growth media [17]. The introduction of cultivation-independent methods has provided a more thorough insight into changes occurring in composition and function of microbial communities [18,19]. This study was designed to investigate the PGP traits and taxonomic composition of soil bacterial community from agricultural fields receiving DE irrigation from more than a decade. For assessing PGP traits, phosphate solubilization, siderophore, indolic compounds and ammonia production were measured for bacterial isolates, and taxonomic characterization of bacterial communities was performed using 16S rRNA gene amplicon sequencing of DNA extracted from bacterial isolates and of DNA directly extracted from soil.

## Materials and methods

### Sampling site and soil collection

The agricultural fields around Gajraula, Uttar Pradesh, India were selected as sampling site, which are being irrigated with DE released from secondary treatment plant of Jubilant Organosys distillery industry for more than a decade. The chemical composition of DE used for irrigation is provided in Table 1. In March 2010, soil samples were randomly collected from a depth of approximately 15 cm using a soil auger along zigzag paths (Zigzag sampling). The samples were transported to the laboratory in an ice-box at 4 °C, and there stored at −20 °C for microbiological analyses. The soil was sandy loam in texture with pH 7.64; electrical conductivity (EC), 0.37 dSm^−1^; organic carbon (OC), 0.64%; total N, 0.08%; Olsen P, 15.86 Kg ha^−1^, extractable K, 367.62 Kg ha^−1^ and, Na^+^, 132 Kg ha^−1^.

**Table 1.**
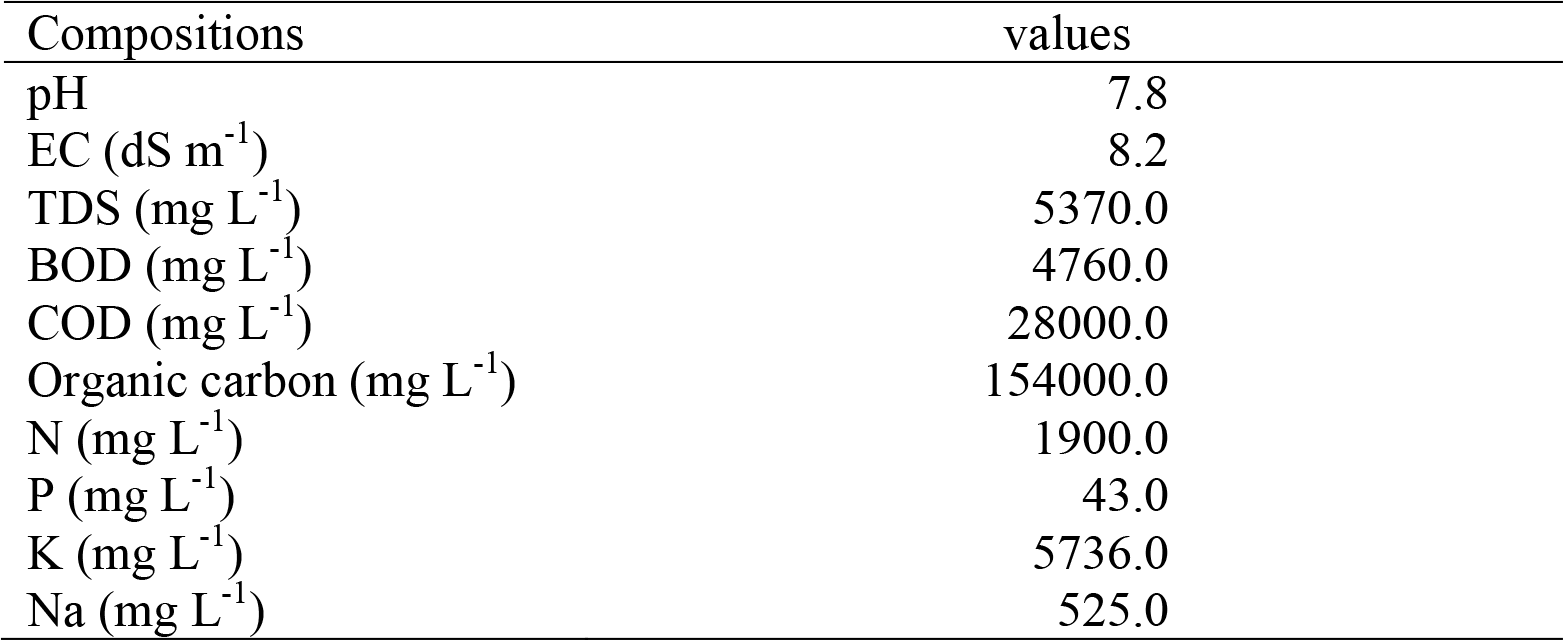
Chemical nature of DE used for irrigation.

### Cultivation of bacterial isolates

Triplicate soil samples (1 g) were taken from each core subsample, homogenized in 10 ml of 0.85% saline, and serially diluted (10-fold dilution) in the same saline. Aliquots (100 μl) were spread on tryptic soy broth agar medium (TSBA; Difco, USA) and Reasoner’s 2A medium (R2A; Difco, USA) plates. Based on morphological differences like shape, size, colour and margin, single colonies were picked at random from the culture plates, purified and maintained on the respective media for further analyses.

### *In vitro* screening of bacterial isolates for PGP traits

All the bacterial isolates were screened for phosphate solubilization on Pikovskaya’s agar plates [20], siderophore production on chrome-Azurol-S agar medium [21], indolic compounds production by Salkowski colorimetric method (indole-3-acetic acid, indole pyruvic acid and indole acetamide) [22] and ammonia production in peptone water [23].

### Taxonomic identification of bacterial isolates

Genomic DNA was extracted from all the bacterial isolates using the method described by Pospiech and Neumann [24], and stored at −20 °C prior to PCR amplification. The 16S rRNA gene was PCR amplified using universal primer pair pA (5′-AGAGTTTGATCCTGGCTCAG-3′) and pH (5′-AAGGAGGTGATCCAGCCGCA-3′) by following the conditions described by Edwards et al. [25]. Approximately 1 μg of PCR products were restricted with endonucleases *Dde* I, and *Taq* I (Fermentas, USA) separately at 37 °C for overnight. The restricted PCR products were resolved by electrophoresis in 2.5% agarose gels and banding pattern was visualized in a gel documentation and analysis system (Alphaimager, USA) using ethidium bromide staining. Strong DNA bands were scored for similarity and clustering analysis using NTSYS pc2.0 program (Applied Biostatistics Inc., USA). The purified PCR products (16S rRNA gene) of representative isolates from each dendrogram cluster was used as template in cycle sequencing reactions using both pA and pH primers with fluorescent dye-labelled terminators (Applied Biosystems, USA). The sequencing was performed in a 3130xl Genetic Analyzer (Applied Biosystems, USA).

The resulting sequences were compared with 16S rRNA gene sequences available in the NCBI GenBank database by BLASTn search (https://blast.ncbi.nlm.nih.gov). The 16S rRNA gene sequences was aligned using CLUSTALW algorithm implemented in MEGA 6.0 [26], and a phylogenetic tree was also constructed in MEGA 6.0 using the neighbor-joining method with 1000 bootstraps. The 16S rRNA gene sequences of bacterial isolates were submitted to NCBI GenBank database under accession numbers HM480326 to HM480328, HM480330 to HM480333, HM480335 and HM480337.

### Soil DNA extraction and clone library construction

Soil DNA was extracted (0.25 g of each sample in duplicate) using MoBio Powersoil™ DNA extraction kit (MoBio Laboratories, USA) according to the manufacturer’s instructions. To remove humic acid contaminations, extracted soil DNA was further purified by using Wizard DNA clean up system (Promega, USA). The 16S rRNA gene was PCR amplified from soil DNA using the same universal primer set (pA and pH) and conditions as used for the DNA of bacterial isolates. All PCR products were purified using QIAEX II Gel Extraction Kit (Qiagen, Germany) and ligated into the plasmid vector pCRII-TOPO (Invitrogen, USA) following the manufacturer’s instructions. The plasmid vectors containing the 16S rRNA gene fragments were transformed into electrocompetent *Escherichia coli* TOP10 cells (Invitrogen, USA). Standard blue/white selection method was used to screen positive clones and checked for right insert size by PCR. All the positive PCR products were restricted with endonucleases *Dde*I and *Hha*I, and restricted PCR fragments were resolved by electrophoresis on 2.5% agarose gels.

### Sequence processing

Representatives of each unique restriction pattern of 16S rRNA gene clones were sequenced on 3130xl Genetic Analyzer (Applied Biosystems, USA) using vector specific M13 forward and reverse primers. All the resulting sequences were analysed using mothur program [27]. First, a set of unique sequences was generated by binning identical 16S rRNA gene sequences. Next, the chimeric sequences were removed using the mothur implementation of UCHIME algorithm [28] in denovo mode. Finally, the sequences were clustered into operational taxonomic units (OTUs) at a threshold of ≥97% sequence similarity using the average neighbor clustering algorithm [29]. Phylogenetic analysis of representatives of each OTU was performed in similar manner as described for bacterial isolates. Representative sequences of each OTU were submitted to GenBank under accession numbers HQ450123 to HQ450150.

## Results and discussion

### Cultivation of bacterial isolates and analysis of PGP traits

A total of 87 bacterial isolates were obtained on TSBA and R2A media. Of these, 45 bacterial isolates were found positive for the PGP traits, including phosphate solubilization (57.3% isolates), ammonia production (54.0% isolates), indolic compounds production (52.4% isolates) and siderophore production (47.5% isolates) (Table 2). Phosphorus availability in soil is vital for growth and development of plants. Phosphate-solubilizing bacteria increase phosphorus availability in soil through solubilization and mineralization of inorganic phosphates, such as Ca_3_(PO_4_)_2_, AlPO_4_, and FePO_4_ [30,31,32]. A high percentage of bacterial isolates recovered form DE irrigated soils showed phosphate solubilization activity, which indicates that DE irrigation did not suppress the growth of phosphate-solubilizing bacterial community. An increase in population size of phosphate-solubilizing bacterial is also reported earlier in agricultural soils irrigated with textile effluent [33]. We have also observed a higher proportion of bacterial isolates positive for ammonia, siderophore and indolic compounds production. Similar to our results, Tripathi et al. [15] found a higher fraction of *Streptomyces* isolates displaying siderophore and indolic compounds production in pulp and paper mill effluent irrigated agricultural soils. Microorganisms produce siderophore to chelate various metals that could be inhibitory to their growth [34], and it has also been reported that production of siderophore enhances production of indolic compounds [35], which possibly explains why we recovered a high percentage of bacterial isolates showing production of both siderophore and indolic compounds.

**Table 2.** Phylogenetic affiliations of representative bacterial isolates.

| Representative isolate | 16S rRNA sequence homology |  |  |
| --- | --- | --- | --- |
|  | Species identified | NCBI accession number | % homology |
| WIFD5 | *Bacillus subtilis* | HM480326 | 99 |
| WIFD12 | *Bacillus megaterium* | HM480327 | 97 |
| WIFD13 | *Paenibacillus pabuli* | HM480328 | 98 |
| WIFD20 | *Bacillus thuringiensis.* | HM480330 | 98 |
| WIFD26 | *Sinorhizobium fredii* | HM480331 | 99 |
| WIFD28 | *Bacillus simplex* | HM480332 | 99 |
| WIFD31 | *Mitsuaria chitosanitabida* | HM480333 | 98 |
| WIFD46 | *Lysobacter yangpyeongensis* | HM480335 | 97 |
| WIFD49 | *Arthrobacter crystallopoites* | HM480337 | 97 |

### Culturable bacterial community

Cluster analysis based on 16S rRNA gene restriction pattern, grouped bacterial isolates into nine clusters. The 16S rRNA gene sequences of one representative isolate from each cluster were identified as *Bacillus megaterium*, *Bacillus simplex, Bacillus thuingiensis, Bacillus subtilis*, *Paenibacillus pabuli*, *Arthrobacter crystallopoites*, *Sinorhizobium fredii*, *Mitsuaria chitosanitabida* and *Lysobacter yangpyeongensis* (Table 3). A phylogenetic reconstruction of 16S rRNA gene sequences of these isolates together with sequences of their nearest relatives is shown in Fig. 1. The collection of bacterial isolates was dominated by *Bacillus* and *Bacillus* derived genera belonged to phylum *Firmicutes* (61%) (Fig. 2a), followed by bacterial genera from Alpha- Beta and Gamma- subdivisions of *Proteobacteria* (31%), and *Actinobacteria* (8%) (Fig. 2a). Members of the phyla *Firmicutes*, *Proteobacteria* and *Actinobacteria* are commonly reported to dominate in culturable soil bacterial communities [16,36,37,38,39].

**Table 3.**
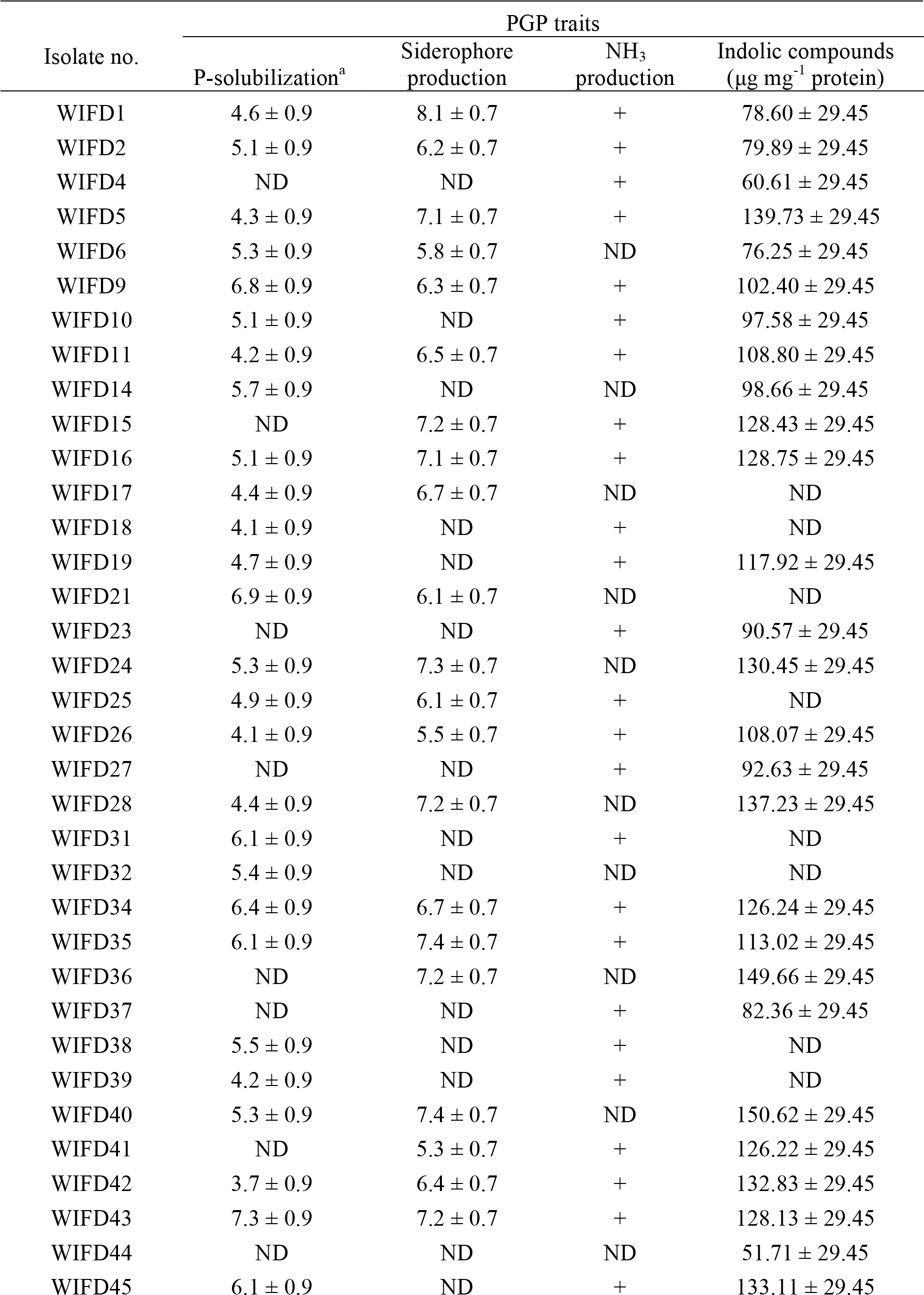
Plant-growth-promoting traits of bacterial isolates cultivated from DE irrigated soil.

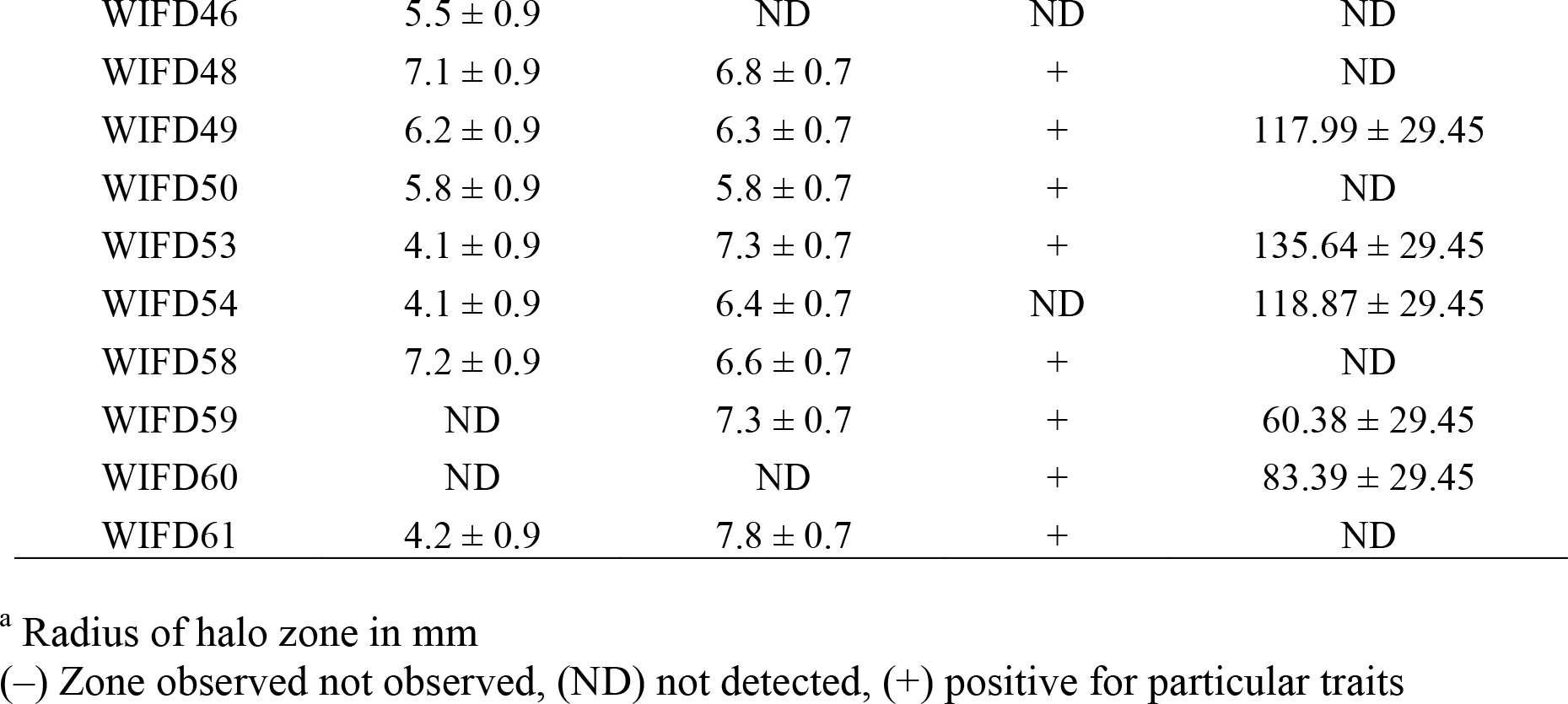

**Fig. 1.**
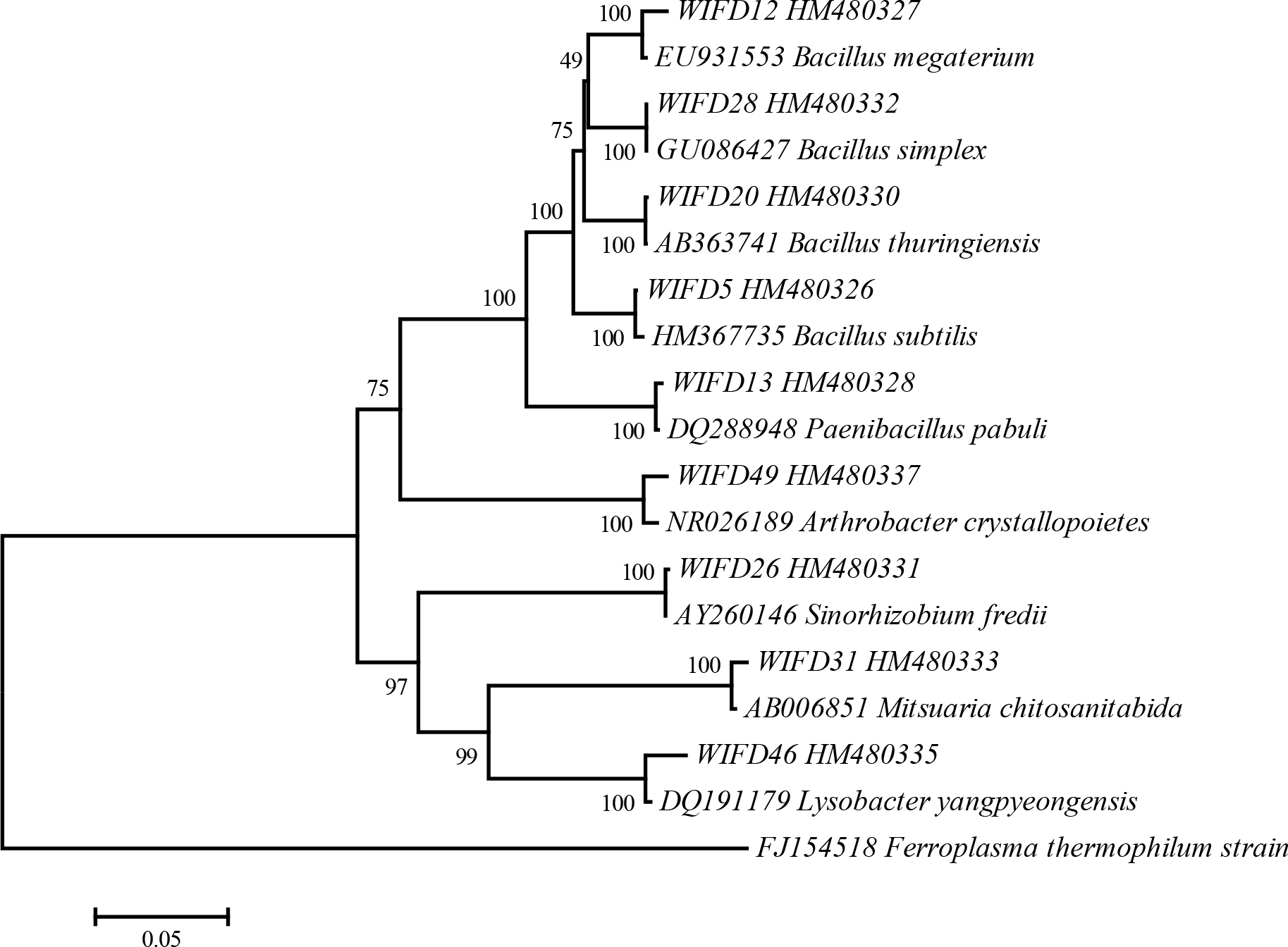
Phylogenetic tree based on the 16S rRNA gene sequences of bacterial isolates cultivated from DE irrigated agricultural soil. The tree was created by the neighbour-joining method. The numbers on the tree indicate the percentage of bootstrap sampling derived from 1000 replicates. 16S rRNA gene sequence of *Ferroplasma thermophilum* used as an out-group.

**Fig. 2.**
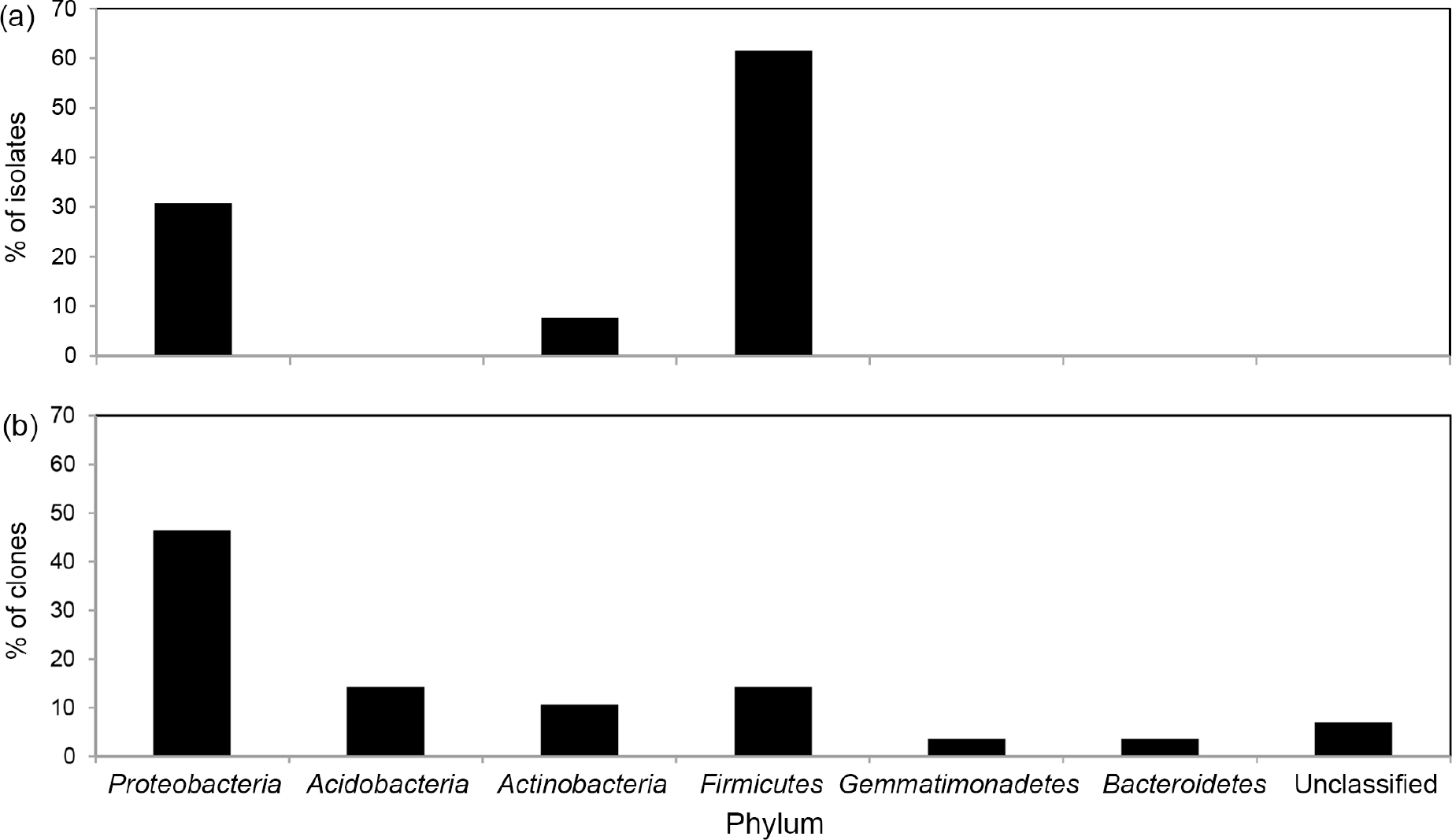
Relative abundance of bacterial phyla in (a) culturable bacterial community and (b) 16S rDNA gene clone library.

### Bacterial 16S rRNA gene clone library

A total of 107 bacterial 16S rRNA gene clones were picked, and after restriction analysis these were grouped in to 57 different clusters. Sequence analysis of these 57 clones indicated presence of 6 chimeric 16S rRNA gene sequences, which were removed prior to further analysis. Chimeric sequences are generated in PCR amplification of environmental DNA by sequence hybridization between closely related microbial taxa [40]. A total of 28 OTUs were detected based on ≥97% sequence similarity. These OTUs were affiliated with six bacterial phyla, including *Proteobacteria* (Alpha-, Beta-, Gamma- and Delta- subdivisions) (46%)*, Acidobacteria* (14%), *Firmicutes* (14%), *Actinobacteria* (11%), *Bacteroidetes* (4%) and *Gemmatimonadates* (4%) (Fig. 2b). Phylogenetic analysis of 16S rRNA gene sequences of these OTUs together with sequences of their nearest relatives is shown in Fig. 3. Bacterial OTUs corresponding to the phylum *Proteobacteria* were most abundant in clone library (Fig. 2b). *Proteobacteria* is the most abundant bacterial phylum in soil clone libraries [41], which is known to contain a great level of physiological and metabolic diversity, and play a crucial role in cycling of carbon, nitrogen and sulfur [42]. *Proteobacteria* is also designated as copiotrophic bacterial taxa [43], which are highly responsive to nutrient amendment [44], and their dominance in clone library could be the result of increased nutrient status of DE irrigated soil. The OTUs belonging to *Acidobacteria*, *Firmicutes* and *Actinobacteria* were the other abundant bacterial phyla in clone library (Fig. 2b). *Acidobacteria* is one of the most dominant soil taxa [45,46], however, due to difficulties associated with cultivation of these taxa, very little is known about their physiology and potential functions [47]. Recent comparative genomic studies have suggested that *Acidobacteria* may play an important role in organic matter decomposition [48]. The proportion of *Firmicutes* OTUs was lower in clone library compared to culturable bacterial community (Fig. 2). This discrepancy is also reported earlier [49,50], and there have been several reasons put forth to explain this difference, which include difficulties associated with lysing endospores during soil DNA extraction and bias in PCR amplification [51]. The members of *Actinobacteria* processes cellulolytic activities which enable them to degrade a wide range of soil organic matter [52]. The increase in actinobacterial population has been reported earlier in soils receiving industrial effluent irrigation [15,53].

**Fig. 3.**
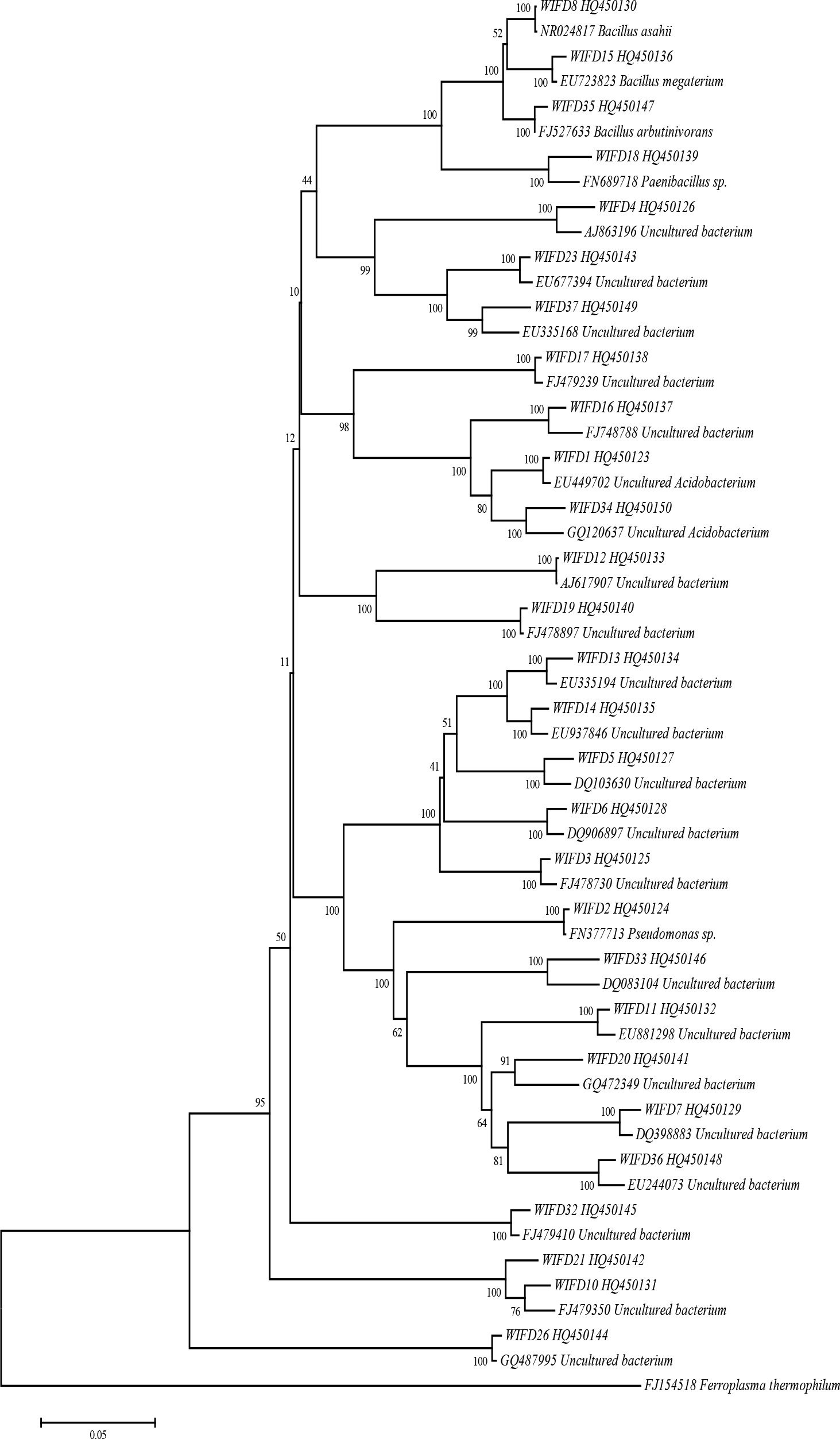
Neighbour-hood joining phylogenetic tree of bacterial OTUs recovered from 16S rRNA gene clone library of DE irrigated soil.

In summary, DE irrigation did not seem to have an adverse effect on PGP traits of culturable bacterial community, therefore using DE in conjugation with irrigation water could be a viable water reuse method in regions facing water scarcity. The cultivation-dependent and - independent methods provided a holistic picture of the bacterial community composition in DE irrigated soil. Further studies focusing on more extensive sampling of DE irrigated agricultural soils in different regions and times of year are necessary to gain better understanding of structure and function of microbial communities.

## Acknowledgements

This work was carried out under network project on Application of Microorganisms in Agriculture and Allied Sectors funded by Indian Council of Agricultural Research, New Delhi. It is certified that there is no financial/commercial conflicts of interest.

